# Four-species *Aspergillus* pan-GWAS reveals rare genome expansion in pathogenicity and contraction in domestication

**DOI:** 10.64898/2026.08.20.745736

**Authors:** Minji Kim, Omid Ardalani, Eduard J. Kerkhoven, Patrick V. Phaneuf

## Abstract

*Aspergillus* species are ecologically diverse and deeply entangled with human health and industry. *A. fumigatus* and *A. flavus* are the two principal species of invasive aspergillosis [1]. *A. niger* and *A. oryzae*, on the other hand, are responsible for global enzyme production, organic acid production [2], and koji-based fermentation industries [3]. The question of whether these similar phenotypes share the same genomic mechanisms across the genus is not yet understood.

To address this, we constructed per-species pangenomes for the four *Aspergillus* species (929 initial genomes filtered to 210 ANI-verified, high-quality assemblies for a total of 88 *A. fumigatus*, 70 *A. flavus*, 33 *A. oryzae*, and 19 *A. niger* assemblies) alongside a genus-level pangenome of 15,163 orthogroups, and conducted phenotype-labeled pan-genome-wide association studies (pan-GWAS) with kinship correction across all species.

Pan-GWAS identified up to 117 significant orthogroup presence/absence associations per species-phenotype comparison. However, convergence analysis showed that among the 92 and 62 distinct gene families significant for human pathogenicity in *A. fumigatus* and *A. flavus* respectively, the two species seldom agreed on whether the pathogenicity was associated with the enrichment or the depletion of a specific gene family. Convergence analysis of the functional annotations also yielded zero significant results at FDR < 0.05. A literature-curated gene panel analysis also showed that a species labeled pathogenic and another labeled GRAS carried the same aflatoxin and virulence genes, suggesting that gene presence alone cannot readily explain their phenotypic differences.

Instead, we propose that niche adaptation operates through the use of the pangenomic rare genome. Reclassifying rare genes by homology identified truly rare subsets (156 to 391 orthogroups per species) distinct from paralogs and gene fragments. Human-pathogenic strains showed significant rare genome expansion of 2.44-fold for both *A. fumigatus* and *A. flavus* (kinship corrected, p = 6.6 × 10⁻⁸). Conversely, industrial strains showed rare genome contraction where both *A. niger* and *A. oryzae* industrial strains carried 0.57-fold (kinship corrected, p = 0.015) fewer rare genes than their non-industrial counterparts.

Hence, we claim that *Aspergillus* niche evolution proceeds through directional rare genome changes, where there is expansion under pathogenic selection, and contraction under industrial domestication. The rare genome, often discarded as noise, may represent the primary evolutionary source for clinical and biotechnological adaptation in this genus.

## 1. INTRODUCTION

Few fungal genera span human health and industry as broadly as *Aspergillus*, with species that range from pathogenic to important producers of fermented foods, enzymes, and pharmaceuticals. *A. fumigatus* is the leading cause of invasive aspergillosis, specified on the World Health Organization fungal priority pathogens list as a critical-priority human pathogen [4]. Similarly detrimental, *A. flavus* is both a human opportunistic pathogen and the primary source of aflatoxin contamination in food crops [5]. Remarkably, the same genus also contains *A. niger* and *A. oryzae*, two important species for industrial biotechnology. *A. niger* is the producer of 80-90% of the world’s citric acid [6], and *A. oryzae* has played an important role in East Asian fermented foods production for millennia [7]. The genomes of *Aspergillus* fungi in these niche environments have been studied extensively within specific species, including virulence-gene sets for *A. fumigatus* [8], [9], [10], [11], mycotoxin and aflatoxin clusters of *A. flavus* [12], [13], [14], and secondary-metabolite repertoires for *A. niger* and *A. oryzae* [3], [15], [16], [17]. However, whether the same genomic architecture, for example a shared “pathogenicity toolkit” or a shared “domestication signature”, persists across the genus has not been tested.

Pangenomic methods can be applied to address this question, through capturing the full gene catalogue of a species rather than a single reference genome. In bacteria, pangenomics has revealed that accessory genomes encode the machinery for niche adaptation, antibiotic resistance, and virulence [18], [19], [20]. In fungi, per-species pangenome studies have addressed the genomic elements of lifestyle variation within individual *Aspergillus* species, identifying biosynthetic gene clusters (BGCs) associated with industrial traits and virulence factors enriched in clinical isolates [9], [21], [22]. However, by design, single-species studies cannot address the question of whether the same lifestyle, when it evolves independently in different species, uses the same genes.

Answering this question requires a multi-species pangenome framework that simultaneously handles pangenome construction, population structure correction, phenotype-labeled genome-wide association testing (pan-GWAS). Comparing those results across species requires a further layer of a genus-level pangenome that provides a common coordinate system for mapping species-specific results.

This study tests whether *A. fumigatus* and *A. flavus*, two pathogenic *Aspergillus* species, converge on the same virulence toolkit, and if not, whether the core, accessory, or rare genome explains this species-specificity. Two economically important domesticates, *A. niger* and *A. oryzae*, will also be tested for the same industrially relevant traits.

We assembled a dataset of 929 *Aspergillus* genomes spanning four species, applied rigorous quality control to retain 210 high-quality assemblies, and constructed per-species pangenomes and a combined genus-level pangenome of 15,163 orthogroups (OGs). We performed phenotype-labeled pan-GWAS with kinship-based population structure correction for each species to preserve the resolution of single-species pangenomes. We then mapped those results onto a genus-level pangenome containing all four species for cross-species convergence analysis. We also identified literature-based genes for phenotypes in the core genome. Ultimately, we propose the rare genome to be where species-specific lifestyle variation is concentrated. The full pipeline and analysis workflow is represented in Figure 1.

**Figure 1:**
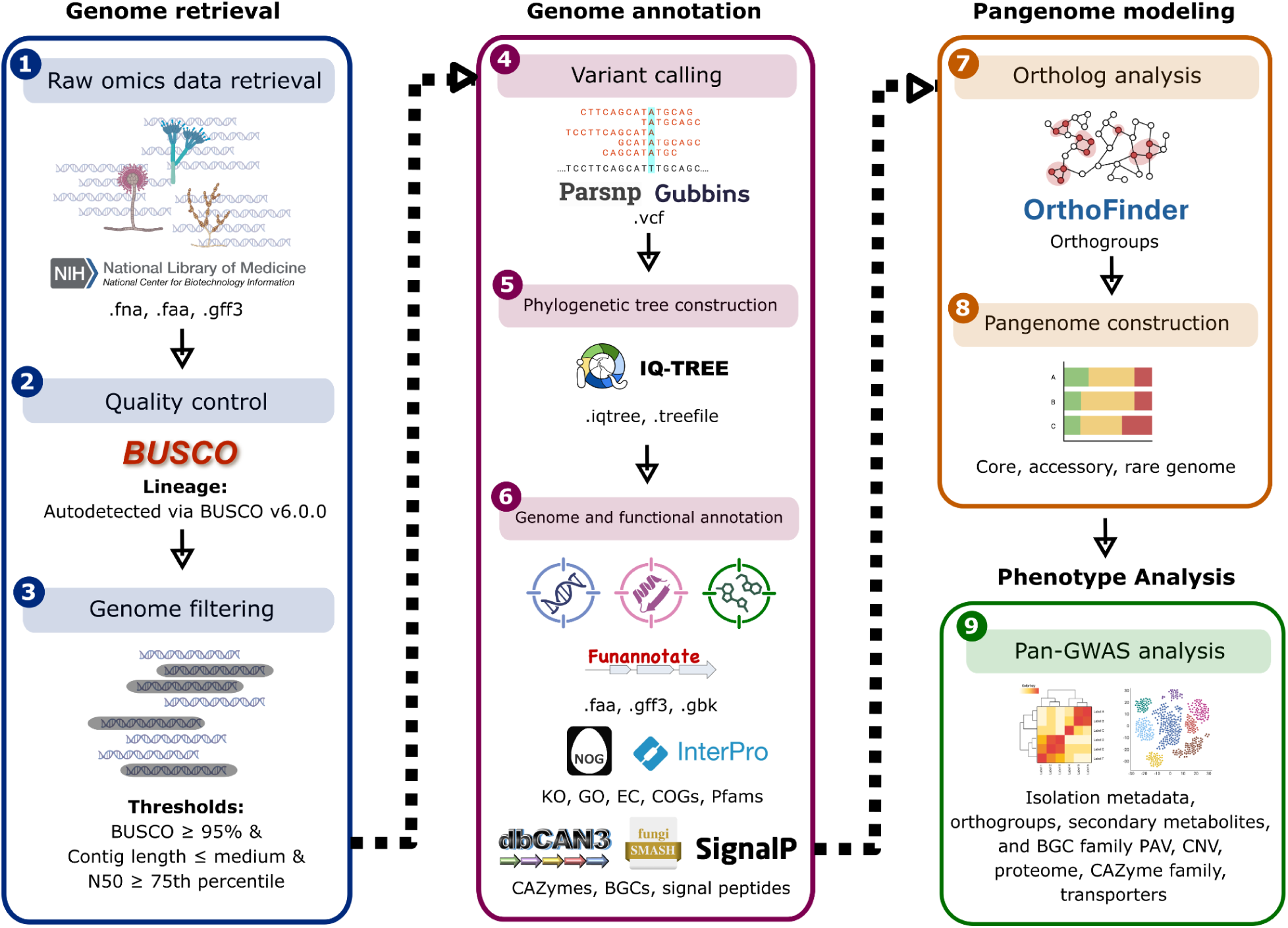
Workflow of pangenome pipeline and data analysis

## 2. METHODS

### 2.1 Genome collection and quality control

We retrieved 929 *Aspergillus* genome assemblies from NCBI [23] spanning four species: *A. fumigatus*, *A. flavus*, *A. niger,* and *A. oryzae*. Quality control was performed using dynamic, species-specific thresholds: BUSCO v6.0.0 [24] completeness score > 95%, contig count ≤ species-specific median, and contig N50 values ≥ species-specific 75th percentile. These dynamic thresholds were selected to account for the variation in sequencing technologies and assembly methods across fungal genomes, which are more heterogeneous than bacterial datasets and therefore not suited to a single global cutoff. After QC, average-nucleotide-identity (ANI) verification was done against the species’ GCF reference at a 95% threshold using fastANI v1.34 [25] and Mash v2.3 [26]. Genomes marked as contaminated from FCS-GX, applied at NCBI submission, were removed from analysis. 210 high-quality assemblies remained: 88 *A. fumigatus*, 70 *A. flavus*, 33 *A. oryzae*, and 19 *A. niger* genomes.

### 2.2 Genome and functional annotation

The 210 assemblies were annotated *de novo* through a unified pipeline. Funannotate v1.8.17 [27] was used for repeat masking (RepeatModeler v2.0.7 [28], RepeatMasker [29]), gene prediction (Augustus [30], GeneMark-ES [31], EvidenceModeler [32]) and integration of protein-evidence and RNA-seq evidence where available. Functional annotation was done using EggNOG-mapper v2.1.13 [33] for COG categories, KEGG orthology, KEGG pathways, Gene Ontology and EC numbers, InterProScan [34] for Pfam, SMART, CDD, PANTHER, SUPERFAMILY and InterPro signatures, SignalP v6 [35] for signal-peptide prediction, and dbCAN3 [36] for CAZyme assignment, including substrate annotation.

### 2.3 Biosynthetic gene cluster analysis

Biosynthetic gene clusters (BGCs) were predicted for all genomes using fungiSMASH v8.0.2 [37]. BGCs were grouped into gene cluster families (GCFs) using BiG-SCAPE v1.1 [38] with clustering cutoff 0.3.

### 2.4 Orthology and pangenome construction

Per-species orthology was inferred with OrthoFinder v3.1.0 [39] (DIAMOND search, MAFFT alignment). A combined four-species OrthoFinder run yielded 15,163 cross-species OGs across 210 genomes. Per-species PAV and CNV matrices were constructed from per-species OrthoFinder gene counts. Per-species OG consensus annotation tables were built by combining EggNOG, InterProScan, dbCAN and SignalP per-protein outputs into one row per OG.

Core, accessory and rare boundaries were not fixed at 95% and 5% but were derived per species by the S-curve inflection method [40]. The OG count histogram was fit in log space with a two-power U-curve P(x) = c₁ · x^(-a₁) + c₂ · (N+1-x)^(-a₂). This resulted in a total of A. fumigatus core ≥ 85, rare ≤ 6, A. flavus core ≥ 68, rare ≤ 6, A. niger core ≥ 18, rare ≤ 2, and A. oryzae core ≥ 32, rare ≤ 3 strains. Pangenome openness was assessed by Heap’s law fit (gamma < 0.05 = closed pangenome) using per-genome incremental sampling with permutation confidence intervals.

### 2.5 Phylogenetic analysis

Per-species core-genome alignments were computed with Parsnp v2.1.5 [41] using each per-species GCF reference. Recombination was identified and masked with Gubbins v3.4.1 [42]. Maximum-likelihood phylogenies were inferred with IQ-TREE v2.4.0 [43] with ModelFinder for substitution-model selection and 1,000 ultrafast bootstrap replicates with Bonferroni-NNI correction [44].

### 2.6 Phenotype classification

A two-tier classifier using manually curated regex rules was used to assign each isolate a Provenance label (where the strain was physically obtained) and a Phenotype label (biological-association label for pan-GWAS). These rules included keywords such as “patient” or “biopsy” for Human-pathogenic associations, and “fermentor” and “bioreactor” for Industrial-trait associations. The classifier combined eight metadata fields (isolation source, BioSample description title, BioProject title and description, WGS reference title, strain identifier, comment, submitter, species name) using a weighted keyword search system. Phenotypes were classified into six categories: Human-pathogenic, Animal-pathogenic, Plant-pathogenic, Industrial-trait, Environmental, and Unknown. For pan-GWAS analyses, Unknown isolates were excluded. Per-genome provenance and phenotype assignments, together with the assembly statistics, accessions and the metadata field and matched keyword that produced each label, are given in Table S1.

### 2.7 Pan-GWAS

Linear-mixed-model EMMA-style [45] association testing was performed per species, per contrast, per feature layer (PAV, CNV, BGC PAV, GCF PAV, GCF CNV). For each feature x, the model y = μ + βx + u + ε was fit, where y is the binary phenotype (case = 1, control = 0), x is the feature, β the fixed-effect coefficient tested against zero, u ∼ N(0, σ²·K) the polygenic random effect with covariance proportional to the kinship matrix K, and ε residual noise. The kinship matrix was computed per species from SNP data and absorbs background relatedness. Power analysis computed minimum-detectable odds ratios at 80% power per contrast under the case-control imbalance.

Pan-GWAS was conducted and false discovery rate (FDR) corrected for each OG association to identify which OGs correlate with phenotypes. Functional enrichment of that significant set was then computed by Fisher’s exact test to see whether the OGs carrying the signal are over-represented in any functional category (CAZy, KEGG, GO, Pfam, COG, EC, InterPro, KEGG_TC, dbCAN substrate). Across every enrichment and convergence tests, uninformative annotation terms were excluded from testing. These included COG-S “Function unknown” and COG-R “General function prediction only”, Pfam terms that start with “DUF”, three GO root nodes (GO:0003674, GO:0008150, GO:0005575), the EC placeholder “-.-.-.-”, and any KEGG_ko/ interpro_IPR/ Pfam terms where the description matches “uncharacterized”, “hypothetical protein”, “protein of unknown function” or “domain of unknown function”.

### 2.8 Cross-species convergence analysis

Per-species pan-GWAS results were mapped into the genus-level pangenome system via protein IDs. In the process, 108 A. fumigatus and 152 A. flavus OGs were added and 326 and 641 OGs were removed due to OG splits and merges, respectively, in the genus-level pangenome. This resulted in 1,655 OGs associated with A. fumigatus and 2,522 OGs associated with A. flavus in the genus-level pangenome. Convergence analysis was done via finding PAV and CNV variants significant in multiple species, and classifying these overlapping variants as concordant (consistent effect directions) or discordant (opposing effect directions). Statistical significance was validated using 1,000-permutation null models that randomly sample OGs from tested pools. Functional convergence was assessed using Jaccard similarity with permutation-based null distributions, followed by testing of individual functional terms using Fisher’s combined probability method with directional concordance effects. FDR correction was applied across annotation and analytical layers. Genus-level pangenome composition was directly compared by summarizing OG counts per functional family (CAZymes, proteases, secreted proteins) across all four species.

### 2.9 Trait-associated gene panel assembly and cross-species comparison

Identification and convergence analysis of recognized trait-associated genes in the Core genome was carried out by curating genes associated with traits from literature, including dedicated genome papers and secondary metabolite cluster databases. A total of 203 *A. fumigatus* virulence genes, 108 *A. flavus* virulence genes, 171 *A. niger* industrial genes, and 127 *A. oryzae* industrial genes were collected. The reference protein of each trait associated with genes were retrieved from NCBI via RefSeq locus tag and a gene-symbol search fallback system, restricted to the target organism. A total of 424 of 609 references were retrieved. Each retrieved protein was BLASTed against our annotated proteome of the reference genome of the four species. 412 of 424 sequences were then successfully linked to the OG IDs in the combined pangenome.

The 412 anchored genes were then filtered in two steps. First, genes were retained only if the source experimental study demonstrated the assigned function at the level of the individual gene and in the correct species, leaving 136 genes. Second, only genes that positively cause their assigned trait were kept. Silenced cluster relics and genes whose deletion increases the trait were excluded. This yielded the final panel of 128 genes (73 virulence, 55 industrial; Table S2). Cross-species conservation was calculated by counting the number of species in which a panel gene was in the core genome, and compared against the rest of the background OGs not in the panel (n = 15,043).

### 2.10 Rare genome characterisation

The rare genome was first reclassified by DIAMOND BLAST [46] against each species’s own Core and Accessory protein pool. The longest representative protein of every Rare OG was queried and the best hits were assigned as either: full-length homolog (≥30 % identity, ≥80 % query and subject coverage; a paralog of a non-rare gene), fragment of longer gene (high subject coverage, low query coverage), no significant similarity (DIAMOND hit below threshold), or no hit (no DIAMOND alignment to Core and Accessory at all). Every Rare OG not in the no-hit class was then excluded, narrowing the Rare genome to a Truly-rare genome.

The truly rare genes were measured on five characteristics: (i) protein length, (ii) Pfam annotation rate, (iii) CAZy annotation rate, (iv) GC-content deviation from genome mean, with a NCBI BLAST of the highest-deviation candidates against nr for cross-kingdom xenolog detection, and (v) per-genome rare OG count by phenotype with Mann-Whitney U testing. COG-category enrichment of the truly rare genome versus Core and Accessory genome was tested by Fisher’s exact test.

### 2.11 Phylogenetic signal and Mash clustering

Fritz & Purvis D phylogenetic signal was computed for every binary phenotype against the recombination-corrected IQ-TREE phylogeny per species, using a 1,000-permutation random null. Mash (sketch size 1,000, k = 21) per-species and combined-species pairwise distances were clustered using both silhouette and elbow criteria for k selection.

## 3. RESULTS

### 3.1 A four-species *Aspergillus* genome resource

To assess whether the assembled resource was clean, verified, and phenotypically balanced enough to support robust cross-species comparison, we retrieved 929 *Aspergillus* genome assemblies from NCBI for the four target species and applied a dynamic threshold system for genome quality filtering. Filters required BUSCO completeness > 95% (autodetected lineage aspergillus_odb12), contig count at or below the per-species median, contig N50 at or above the per-species 75th percentile, and at least one RefSeq (GCF) reference assembly per species. Together with ANI verification against the GCF reference (fastANI v1.34, 95% threshold) per species, 210 final assemblies were left. Of the 210 high quality assemblies, 88 were of *A. fumigatus*, 70 *A. flavus*, 19 *A. niger*, and 33 *A. oryzae*. Per-species RefSeq references were GCF_000002655.1 for *A. fumigatus*, GCF_014117465.1 for *A. flavus*, GCF_000002855.4 for *A. niger* and GCF_000184455.2 for *A. oryzae*. Across the filtered strains, BUSCO completeness was high (mean 99.0%, minimum 95.9%), with median contig N50 of 890.7 kb and median total assembly length of 36.5 Mb.

A classifier integrating eight metadata fields (isolation source, BioSample description, BioProject title and description, WGS reference title, strain identifier, comment, submitter and species name) was used to assign each isolate into two layers of labels. The Provenance label named where the strain was physically isolated from (Clinical, Environmental, Industrial-origin, Culture-derived, Unknown), and the Phenotype label specified the biological association used for pan-GWAS (Human-pathogenic, Environmental, Industrial-trait, Plant-pathogenic, Animal-pathogenic, Unknown). Across the 210 genomes the Provenance distribution was 122 Clinical (58.1%), 44 Environmental (21.0%), 26 Industrial-origin (12.4%), 9 Culture-derived (4.3%) and 9 Unknown (4.3%). The same metadata was used to record the collection site for 186 of the 210 genomes (88.6%), whose geographic distribution is portrayed in Figure 2.a. *A. fumigatus* was the most geographically diverse, with isolates from 14 countries in 29 separate BioProjects, representing its role as a worldwide airborne saprophyte. *A. oryzae* was geographically the least diverse, referenced by 20 BioProjects but isolated from just three countries (South Korea, China, and Japan), reflecting its history of use and domestication from the koji mold.

**Figure 2.**
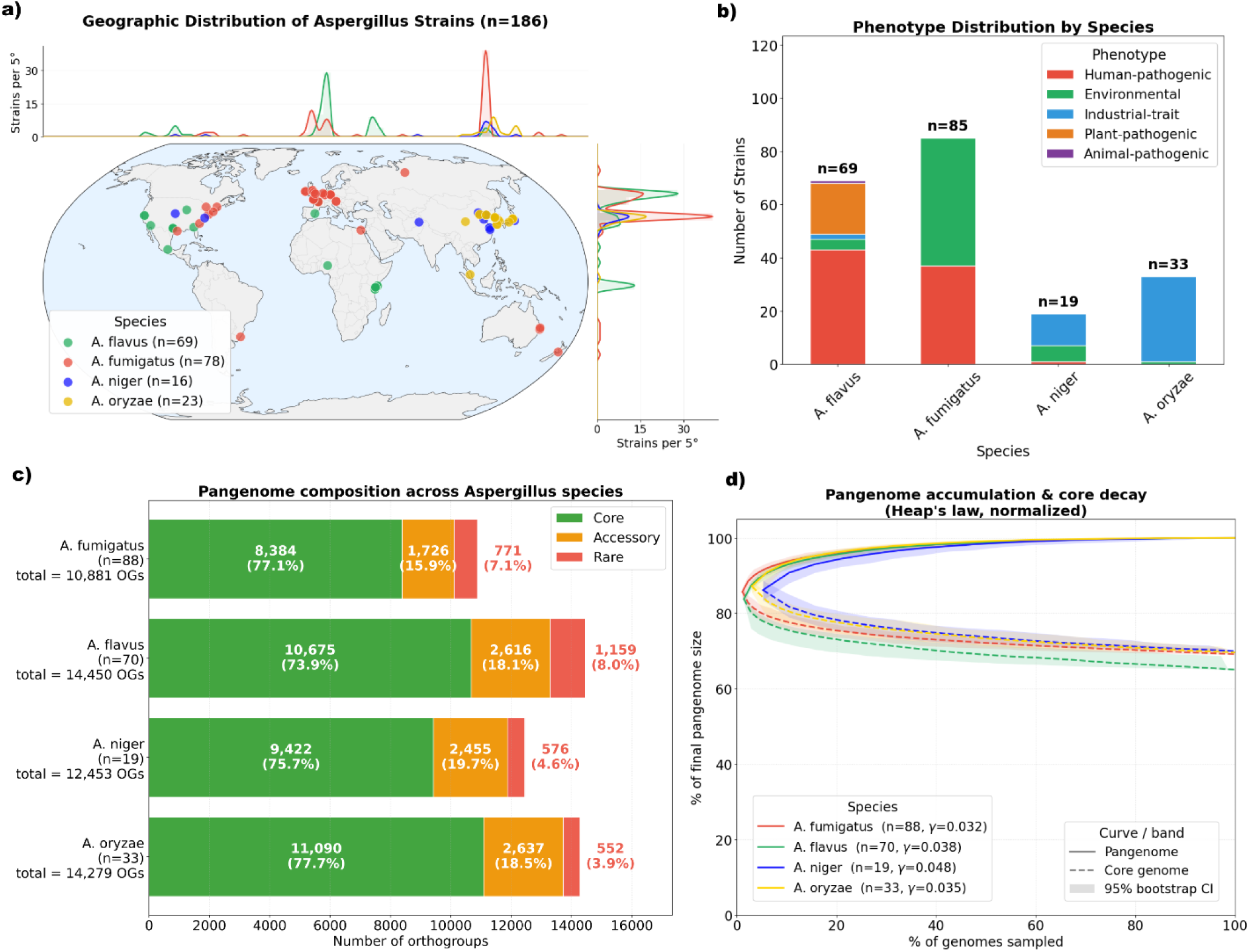
A four-species Aspergillus genome resource (n = 210). a) Geographic distribution of the origin of all 210 quality-filtered, ANI-verified genomes. b) Per-species phenotype distribution after classification. c) Per-species pangenome composition based on the number of orthogroups (OGs) classified as core, accessory, or rare genes. d) Heap’s law pangenome growth curves with 95% permutation confidence ribbons.

The Phenotype distribution was 81 Human-pathogenic (38.6%), 59 Environmental (28.1%), 46 Industrial-trait (21.9%), 19 Plant-pathogenic (9.0%), 1 Animal-pathogenic and 4 Unknown. Excluding the four Unknown-phenotype isolates resulted in a 206-genome analysis dataset. Phenotype distributions, as shown in Figure 2.b, were strongly asymmetric within each species, where *A. fumigatus* (88) strains had 48 Environmental, 37 Human-pathogenic and 3 Unknown strains, *A. flavus* (70) had 43 Human-pathogenic, 19 Plant- pathogenic, 4 Environmental, 2 Industrial-trait, 1 Animal-pathogenic and 1 Unknown, *A. niger* (19) had 12 Industrial-trait, 6 Environmental and 1 Human-pathogenic, and *A. oryzae* (33) had 32 Industrial-trait and 1 Environmental. In *A. oryzae*, due to the drastic phenotype imbalance, we designated its results as informative for only descriptive statistics. Overall, the resource is clean enough to compare the four species directly, but the uneven within-species phenotype splits mean that *A. fumigatus*, *A. flavus* and *A. niger* carry the association testing and *A. oryzae* is retained for descriptive statistics only.

### 3.2 Closed pangenomes across the four species

To establish whether our sampling was saturated, and whether the core, accessory and rare compartments already foreshadow lifestyle signals, OrthoFinder v3.1.0 was run on protein predictions (from Funannotate) of each species independently. For each species we built two OG by genome matrices: a presence-absence variance (PAV) matrix recording each OG as present (1) or absent (0) per genome, and a copy-number variance (CNV) matrix recording its number of gene copies per genome. PAV captures gene gain and loss, revealing traits that depend on the presence of a gene, while CNV captures gene duplications and copy losses, revealing traits that depend on how many copies a gene has. Both matrices were tested as independent pan-GWAS layers. OGs were labelled as Core, Accessory, or Rare genes using per-species gene-frequency thresholds derived by the S-curve inflection-point method. The pangenome composition of each species is shown in Figure 2.c. Heap’s law fits, as shown in Figure 2.d, gave gamma values well below 0.05 in all four species, indicating that the current batch of samples would not gain much new gene content with each additional sequenced isolate. Per-species pangenome statistics are summarised in Table 1.

**Table 1:** Pangenome summary statistics for four Aspergillus species. OG denotes orthogroup.

| Species | Genomes | Total OGs | Core OGs (%) | Accessory OGs (%) | Rare OGs (%) | Heap's $\gamma$ | Pangenome Status |
| --- | --- | --- | --- | --- | --- | --- | --- |
| *A. fumigatus* | 88 | 10,881 | 8,384 (77.1%) | 1,726 (15.9%) | 771 (7.1%) | 0.032 | Closed |
| *A. flavus* | 70 | 14,450 | 10,675 (73.9%) | 2,616 (18.1%) | 1,159 (8.0%) | 0.038 | Closed |
| *A. niger* | 19 | 12,453 | 9,422 (75.7%) | 2,455 (19.7%) | 576 (4.6%) | 0.048 | Closed |
| *A. oryzae* | 33 | 14,279 | 11,090 (77.7%) | 2,637 (18.5%) | 552 (3.9%) | 0.035 | Closed |

Across the four species, functional enrichment analysis of the accessory genome showed a conserved depletion signal but species-specific enrichment. In all four species, the accessory genome was significantly depleted (Fisher’s exact, q < 0.05) of housekeeping functions. COG categories J (translation), A (RNA processing), and U (intracellular trafficking and secretion) were all depleted in the accessory and instead consistently resided in the core genome. On the other hand, ankyrin-repeat Pfam families were significantly enriched in the accessory genome for Ank_4 in *A. flavus* and Ank_2 in *A. niger*. NACHT-domain OGs (NLR-like sensor/effector architecture) were also significantly enriched in *A. niger* and *A. oryzae*. CAZy enrichment in the accessory genome was species-specific. *A. niger* was enriched for chitin-binding molecules (CBM18, CBM50) and chitinase (GH18), *A. oryzae* for glycosyltransferase (GT25) and laccase/multicopper oxidases (AA1_3), and *A. flavus* for α-L-rhamnosidases (GH78). Finally, mycotoxin biosynthesis (GO:0043386) was significantly enriched in the accessory genomes of *A. flavus* and *A. oryzae*.

Several of these enrichments align with established phenotypes of each species. The mycotoxin-biosynthesis (GO:0043386) in *A. flavus* and *A. oryzae* reflects the characterized contrast between *A. flavus*, an aflatoxigenic crop and food contaminant, and its domesticated, Generally Recognized As Safe (GRAS) relative *A. oryzae*. Although the aflatoxin cluster is retained in *A. oryzae*, the species is non-aflatoxigenic, with industrial, non-production strains attributed to disabling mutations [47]. In the same way, the enriched CAZy families of CBM18/CBM50 chitin-binding modules and GH18 chitinases in *A. niger* fit its saprotroph and industrial enzyme-producer role, the GH78 α-L-rhamnosidases in *A. flavus* its colonisation of plant tissue, and the AA1_3 laccases in *A. oryzae* the phenolic oxidation of its koji fermentation.

The accessory genome of the genus is therefore conserved in its depletion of core-housekeeping COGs, while its enriched contents such as CAZy and mycotoxin composition varies across the genus. These enrichments therefore already separates these species by lifestyle, but it does so at the level of the species as a whole. Whether gene content also separates strains of differing phenotype within a species is tested next using pan-GWAS.

### 3.3 Pan-GWAS reveals minimal cross-species shared associations

Having established which functions are enriched at the level of the species, we next asked which specific OGs track phenotype among strains within each species. For every species with at least two phenotype groups, one-vs-rest binary contrasts were constructed. This binary contrast was chosen over a pairwise contrast, so that significant hits could be attributed to a single phenotype. Associations were fitted with an EMMA-style linear mixed-model (LMM) with SNP-derived genetic relatedness (kinship) matrix as random-effect covariance, to reflect true phenotype instead of population structure. Per species, the pan-GWAS tests were run across the same PAV and CNV matrices, subset to phenotypes. Tests were also run on fungiSMASH-derived biosynthetic gene cluster (BGC) PAV matrices and on BiG-SCAPE-derived gene cluster family (GCF) PAV/ CNV matrices. The Benjamini-Hochberg (BH) false-discovery-rate (FDR) threshold was set at q < 0.05. The complete number of significant associations per feature layer is shown in Figure 3.a.

**Figure 3.**
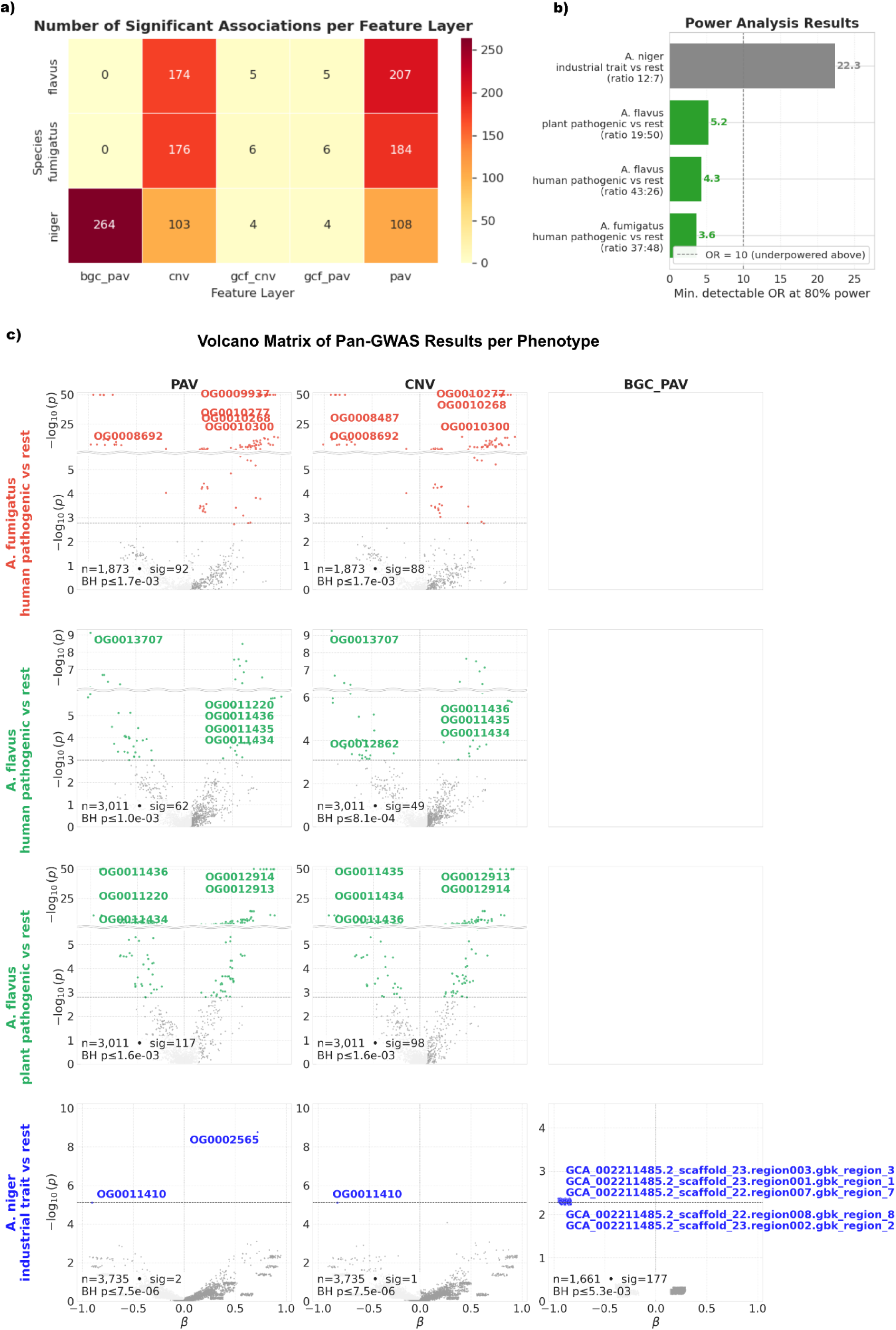
Per-species pan-GWAS results according to species-specific associations. a) Heatmap of total significant hits across all species, comparisons, and feature layers. PAV presence-absence variation, and CNV copy-number variation. BGC denotes a biosynthetic gene cluster (from antiSMASH) and GCF a gene cluster family (from BiG-SCAPE); the BGC_PAV, GCF_PAV, and GCF_CNV columns report PAV hits at the BGC and GCF levels, and CNV at GCF levels, respectively. b) Minimum detectable odds ratio (OR) per species at 80% power per comparison. c) Volcano matrix of EMMA-style linear mixed-model pan-GWAS associations across comparisons and three feature layers (PAV, CNV, BGC_PAV). OG denotes orthogroup. Coloured points pass Benjamini-Hochberg (BH) FDR < 0.05, the dashed line marks the BH cutoff p-value, and the top hits by |β| are labelled per panel.

Isolates labeled Unknown were excluded, as this label reflects missing metadata or undefinable phenotype. After excluding the Unknown phenotype label, the pan-GWAS sample sizes were 85 *A. fumigatus*, 69 *A. flavus* and 19 *A. niger* isolates. *A. oryzae* yielded zero contrasts because all 32 industrial isolates were paired with only one environmental isolate, an imbalance that failed the minimum n_case/ n_control threshold for the LMM. Power analysis showed that the *A. fumigatus* human-vs-rest, *A. flavus* human-vs-rest and *A. flavus* plant-vs-rest contrasts were adequately powered, with minimum detectable odds ratios at 80% power of 3.61, 4.28 and 5.25, respectively. The power analysis results are illustrated in Figure 3.b. The *A. niger* industrial-vs-rest contrast was underpowered, requiring an odds ratio of 22.33 for 80% power; the scarce number of significant hits in *A. niger* therefore cannot be interpreted as evidence of true negatives. The number of significant OG hits per species and contrast are summarised in Table 2.

**Table 2.** Pan-GWAS hits across four Aspergillus species (FDR < 0.05). OG denotes orthogroup, PAV presence-absence variation, and CNV copy-number variation; the OG PAV and OG CNV columns report hits from the OG presence-absence and copy-number layers. BGC denotes a biosynthetic gene cluster (from antiSMASH) and GCF a gene cluster family (from BiG-SCAPE); the BGC PAV and GCF PAV columns report presence-absence hits at the gene cluster and cluster-family levels.

| Species | Contrast | n (case/control) | OG PAV | OG CNV | BGC PAV | GCF PAV |
| --- | --- | --- | --- | --- | --- | --- |
| *A. fumigatus* | Human pathogenic vs rest | 37 / 48 | 92 | 88 | 0 | 3 |
| *A. fumigatus* | Environmental vs rest | 48 / 37 | 92 | 88 | 0 | 3 |
| *A. flavus* | Plant pathogenic vs rest | 19 / 50 | 117 | 98 | 0 | 4 |
| *A. flavus* | Human pathogenic vs rest | 43 / 26 | 62 | 49 | 0 | 1 |
| *A. flavus* | Environmental vs rest | 4 / 65 | 28 | 27 | 0 | 0 |
| *A. niger* | Industrial vs rest | 12 / 7 | 2 | 1 | 177 | 2 |
| *A. niger* | Environmental vs rest | 6 / 13 | 106 | 102 | 87 | 2 |

The volcano matrix in Figure 3.c shows the significant associations concentrated in three feature layers. Half (51.9%) of the significant hits of pan-GWAS hits have generic annotations or are not annotated. Of those that are annotated in A. fumigatus human pathogenic strains, notable hits include ferric reductase (OG0010109), enriched, and MFS sugar transporter (OG0009116), depleted, in human-pathogenic strains. The MFS sugar transporter is one of A. fumigatus’ primary carbon uptake routes from soil and compost as a saprotroph, carried by 98% of the environmental strains, however, the human lung presents a narrow, host-derived nutrient landscape where such a transporter can be dispensable, carried only by 30% of the human-pathogenic strains.

In A. flavus human-pathogenic strains, the MACPF/ perforin domain (OG0011496), a membrane-damaging effector, was enriched. In A. flavus plant-pathogenic strains, a siderophore-iron-reductase-like cluster of four genes (OG0012600 AMP-binding (adenylation/acyl-CoA ligase), OG0012599 amidohydrolase, OG0012598 MFS transporter, and OG0012490 FhuF-like ferric reductase) was enriched, and GH18 chitinase (OG0000295), involved in fungal cell wall remodeling and autolysis of itself and antagonistically toward competitors, was depleted.

In *A. niger*, the only enriched PAV hit for industrial strains is the PDR-family ABC transporter (OG0002565), an efflux pump of drugs and metabolites.

Functional enrichment of the annotated, significant OGs was computed per contrast, for each functional annotation layer, by Fisher’s exact test and FDR correction within each layer. Only two terms reached q < 0.05 across all species and contrasts. *A. flavus* plant-pathogenic strains were enriched for secondary metabolism (COG-Q, q = 0.020). *A. niger* industrial-trait strains were again enriched for the pleiotropic-drug-resistance/ CDR ABC-transporter efflux family (KEGG_TC 3.A.1.205, q = 0.016), carried by OG0002565. *A. fumigatus* human-pathogenic and *A. flavus* human-pathogenic contrasts produced no terms surviving q < 0.05.

Cluster-level BGC and GCF pan-GWAS revealed two distinct patterns. At the BGC level, almost every significant feature has a minor allele frequency (MAF) near 1 / n_strains and represents a single-strain-specific region rather than a cross-strain phenotype signal. This is depicted by 177 significant BGC PAV features in the *A. niger* industrial contrast all at MAF = 0.053 (1/19). The features are dominated by NRPS and NRPS-like clusters (116/177, 66%), with smaller contributions from type I polyketide (29), terpene (26) and indole (6) clusters. At the GCF level the signal is sparser but more interpretable. *A. fumigatus* human-pathogenic-vs-rest yielded three significant GCFs (FAM_00050, FAM_00074, and FAM_00070), *A. flavus* plant-pathogenic-vs-rest yielded four significant GCFs (FAM_00152, FAM_00059, FAM_00077, and FAM_00044), and *A. flavus* human-pathogenic-vs-rest yielded one (FAM_00077) family that is gained in human-pathogenic *A. flavus* strains but lost in the plant-pathogenic strains of the same species. FAM_00077 was the only GCF with a characterized reference compound in MIBiG, encoding diketopiperazine aspirochlorine (MIBiG BGC0001123). *A. niger* industrial vs rest gave 2 significant GCFs (FAM_00036 and FAM_00189), both lost in industrial strains.

Each powered contrast therefore yields interpretable phenotype-tracking gene content, but the associations share no evident common thread by inspection. Convergence is therefore tested formally below, on a shared coordinate system of a genus-level pangenome, rather than asserted from the results above.

### 3.4 Cross-species convergence using the genus-level pangenome

To test whether the two pathogens converge, that is, whether they agree on both the identification and the direction of effect of phenotype-associated gene content, we projected the per-species results onto a common coordinate system that these species otherwise lack. A combined OrthoFinder analysis across the protein predictions of all 210 genomes produced a genus-level pangenome of 15,163 cross-species orthogroups, to which the significant hits from each pan-GWAS were mapped via protein lookup tables. Convergence was then tested at three levels of resolution: individual orthogroups, functional annotation terms, and secondary-metabolite gene-cluster families. At none of the three did A. fumigatus and A. flavus converge.

#### 3.4.1 Orthogroup-level convergence

We first asked whether, among the OGs jointly tested in both species, the directions of effect agree. For the human-pathogenic-vs-rest contrast, *A. fumigatus* and *A. flavus* contributed 1,655 and 2,522 genus-level OGs at the PAV layer, with 448 jointly tested. The cross-species directional convergence is portrayed in Figure 4.a. Of the 448, 221 (49.3%) were directionally concordant (both positive or both negative beta coefficients) and 227 (50.7%) were directionally discordant. Of the 221 concordant OGs, 139 had positive betas in both species (gained in human-pathogenic strains) and 82 had negative betas in both species (lost in human-pathogenic strains). The same analysis on the CNV layer gave 448 jointly tested OGs with 220 (49.1%) concordant and 228 (50.9%) discordant. Industrial-trait convergence between *A. niger* and *A. oryzae* was not estimable as *A. oryzae* generated zero significant OGs under the case/control imbalance constraint. No OG reached significance in both species at either layer, even with a 1,000-permutation null model. The 49.3% (PAV) and 49.1% (CNV) directional-concordance ratios were indistinguishable from the 50:50 expectations (exact binomial p = 0.81 and p = 0.74), and effect sizes were uncorrelated between species (Spearman ρ = −0.023 and −0.020).

**Figure 4.**
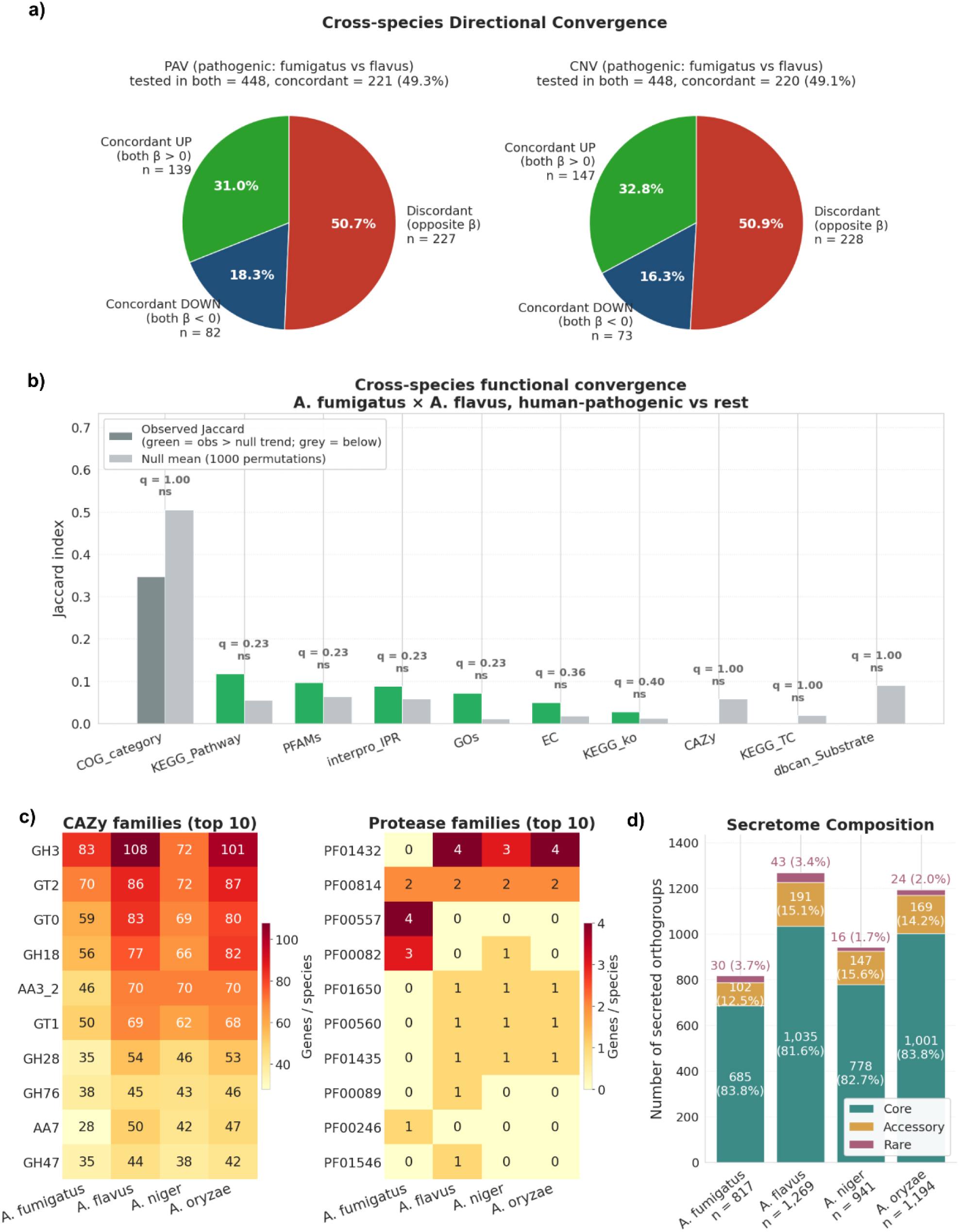
Cross-species convergence is predicted to be absent. a) Directional convergence of orthogroups significant in both A. fumigatus and A. flavus (human-pathogenic vs rest), split by sign of effect. b) Functional convergence per annotation layer shown via a comparison of observed against 1000-permutation null mean of the two species’ enriched sets. Observed bars are coloured green if the observed Jaccard exceeds the null mean (a directional trend toward convergence) and grey otherwise. c) Per-species gene counts for the top 10 CAZy families (left) and top 10 protease families (right). d) Predicted secretome per species via SignalP.

#### 3.4.2 Functional-level convergence

We next asked whether, even without sharing the exact OGs, the two species enrich for the same functions and in the same direction. For every species pair, we computed the Jaccard similarity of the two species’ pan-GWAS-significant OG functional terms and compared it against a 1,000- permutation Jaccard null. As illustrated by the graph in Figure 4.b, 0/ 30 comparisons survived BH-FDR q < 0.05. The strongest individual trends were *A. fumigatus* and *A. flavus* environmental vs rest sets sharing one CAZy family (GH18, p = 0.052) and three KEGG pathways under the human-pathogenic contrast (p = 0.063). But these did not survive multiple-testing corrections. Therefore, functional comparisons did not rescue convergence, in any species pair or contrast.

#### 3.4.3 Gene-cluster-level convergence

We then asked whether the species share GCFs along the secondary-metabolite dimension that OG and functional testing failed to reveal. An analysis of the BiG-SCAPE-derived GCFs returned 0 of 34 GCF families significant across the *A. fumigatus* vs *A. flavus* pathogenic comparison. The pathogenic and industrial gene sets of these species therefore also do not converge at the GCF level. With OG, functions and GCF levels all negative, the absence of convergence is clearly consistent. With the OG, functional and GCF levels all negative, the absence of convergence is consistent across every resolution tested.

#### 3.4.4 Functional family annotation

Independent of pan-GWAS significance, we asked whether the four pangenomes are built from similar raw inventories of CAZymes, proteases and secreted proteins, within which lifestyle differences could reside. We calculated per-species OG counts in each CAZy, protease and secretome family directly from the OG consensus tables. 273 CAZy families were detected across the four species’s pangenomes. The top 10 CAZy families detected are shown in Figure 4.c (left). The most expanded CAZy families consist of fungal cell wall biosynthesis and plant cell wall hydrolysis, while appearing in all four species at proportionally similar OG counts. The number of predicted protease OGs, mainly made up of housekeeping peptidase families, are shown in Figure 4.c (right). Secretome size was 817 (7.5%) in *A. fumigatus*, 1,269 (8.8%) in *A. flavus*, 941 (7.6%) in *A. niger* and 1,194 (8.4%) in *A. oryzae*. As illustrated in Figure 4.d, the vast majority of secreted OGs were Core in every species (685 to 1,035), with only 16 to 43 in the Rare genome.

The four species are therefore built from proportionally similar functional inventories, so lifestyle differences cannot be attributed to the CAZyme, protease or secretome repertoires. Pan-GWAS and cross-species convergence have so far tested mainly the variable, accessory genome, and our results have shown lifestyle cannot be attributed to it. Hence, the core genome, which holds most of the OGs in all four species, is examined next.

### 3.5 Literature-curated virulence and industrial genes are conserved across the core genome of all species

As pan-GWAS is by construction restricted to variable gene content, we asked whether the OGs responsible for lifestyle are invisible to the association tests because they sit in the core genome, and whether core gene content can itself explain species-specific lifestyle. A curated literature search was therefore conducted to identify trait-associated genes. Out of 412 genes retrieved that could be anchored to our OGs via a BLAST search against each species’ proteome, a final panel of 128 (73 virulence, 55 industrial) experimentally validated, positively associated genes were kept (Table S1), corresponding to 120 unique OGs.

As portrayed in Figure 5.a, 108 of 128 trait-associated genes (84%) are core and conserved in all four species, compared with the 41% in the background OGs (Mann-Whitney, p = 2 x 10⁻²²). None of the 73 virulence genes were pathogen-specific, that is, core in *A. fumigatus* and *A. flavus* and absent in both *A. niger* and *A. oryzae*. As for the 20 genes that are variable (17 core in some, 3 not core in all four species), these were concentrated in *A. fumigatus* secondary metabolite and stress genes (the gliotoxin cluster, zinc-homeostasis genes), *A. niger* industrial enzymes and transporters (phytases, sugar transporters, glucose oxidase), and the *A. oryzae* kojic-acid locus (Figure 5.b (top)).

**Figure 5.**
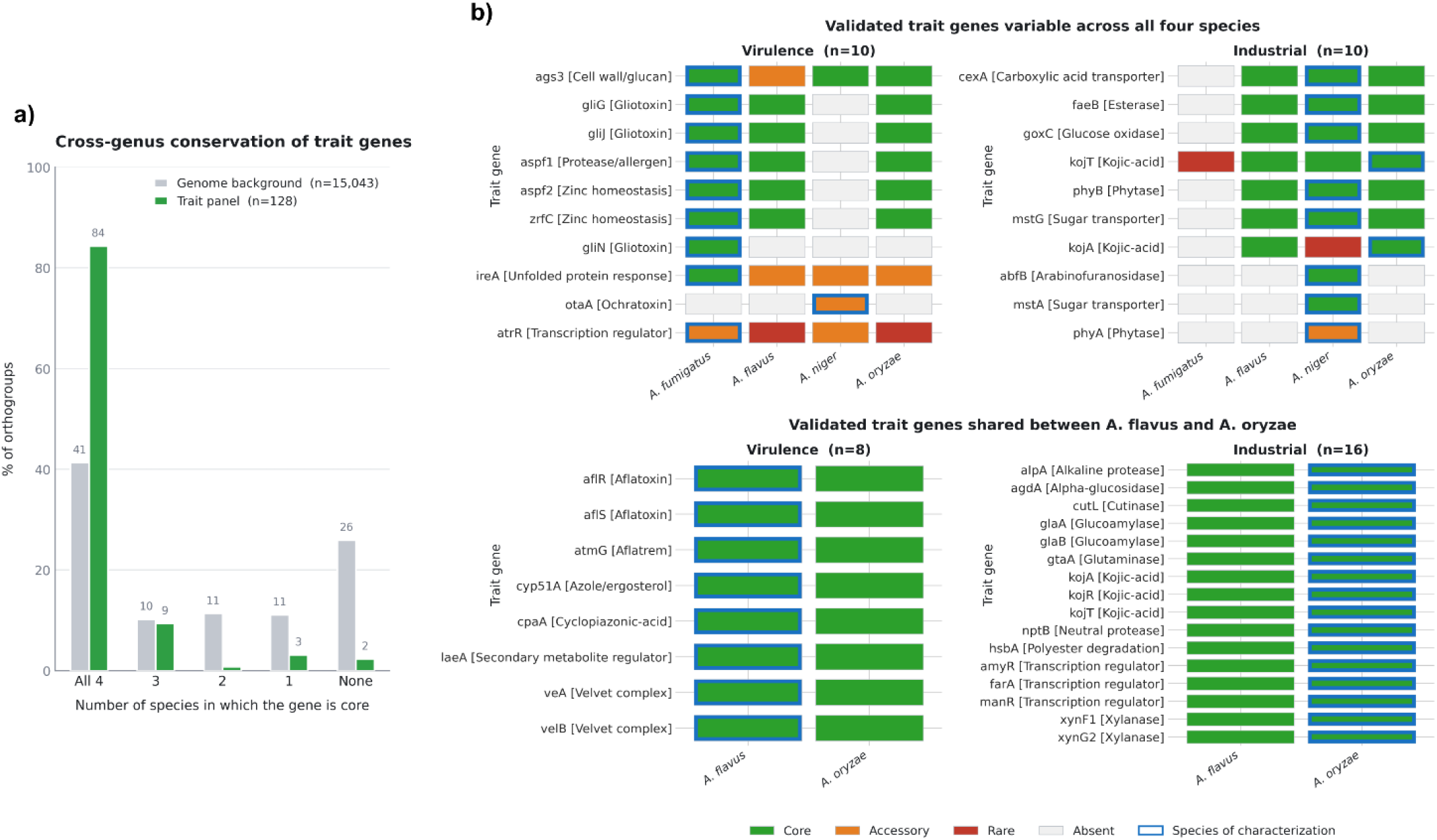
Literature-curated virulence and industrial genes are conserved across the core genome. a) Percentage of orthogroups core in species for the background (grey, n = 15,043) and the curated (green, n = 128) genes. b) Pangenome class (core, accessory, rare, or absent) of each validated trait gene. The blue outline marks the species in which each gene was experimentally characterized for its trait in literature. Top: genes that are not core in all four species, shown across all four species, with genes associated with virulence (left, n = 10) and industrial (right, n = 10). Bottom: genes shared between A. flavus and A. oryzae, shown for A. flavus virulence (left, n = 8) and A. oryzae industrial (right, n = 16) genes.

The *A. flavus* and *A. oryzae* pair shows that this conservation holds even between species of opposing phenotypes. *A. oryzae* is largely understood as the GRAS, food-grade, non-toxigenic domesticate of the aflatoxigenic *A. flavus* [48], yet all 8 experimentally validated, core, virulence-associated *A. flavus* genes are also core in *A. oryzae*. Symmetrically, all 16 validated, core, industrial-associated *A. oryzae* genes are also core in *A. flavus* (Figure 5.b). The two species therefore essentially overlap in virulence and industrial gene set.

In summary, canonical trait genes are indeed largely invisible to pan-GWAS because 84% are core in all four species, and that same conservation disqualifies them as the explanation for lifestyle, as the genes are shared even by a critical-priority pathogen and a GRAS domesticate.

### 3.6 Reclassification of the rare genome reveals bidirectional, phenotype-associated size change

To determine whether the rare genome, once purified of noise, carries a reproducible and directional signal separating the two pathogens from the two domesticates, we first partitioned it by homology. Frequency alone groups the rare genome into three biologically distinct populations: paralog splits of non-rare gene families, fragmented genes that align partially with longer non-rare proteins, and genuinely strain-exclusive within the species’ OGs with no internal homologs. To separate these populations, we DIAMOND BLASTed every rare OG’s representative protein against its own species’s core and accessory protein pool. Each rare OG was categorized into: Full-length homolog (paralog of a non-rare gene; ≥ 30% identity, ≥ 80% query and subject coverage), Fragment of longer gene (high subject coverage, low query coverage), No significant similarity (DIAMOND hit below threshold), or No hit (no DIAMOND alignment to Core and Accessory at all). Across the four species, the Full-length-homolog (paralog) class accounted for 13.4% to 14.2%, the Fragmented class for 20.5% to 40.9%, the No-significant-similarity class for ∼17 %, and the No-hit (truly rare) class for 28.3% to 47.9% of the original rare OGs. The results of the DIAMOND classification per species is shown in Figure 6.a with per-OG hit statistics, coverage values and pangenome core, accessory, and rare genome assignments given in Table S3. We filtered out every Rare OG not in the No-hit class and restricted all of the following rare-genome analyses to keep the truly rare, strain-exclusive within the species’ subset only.

**Figure 6.**
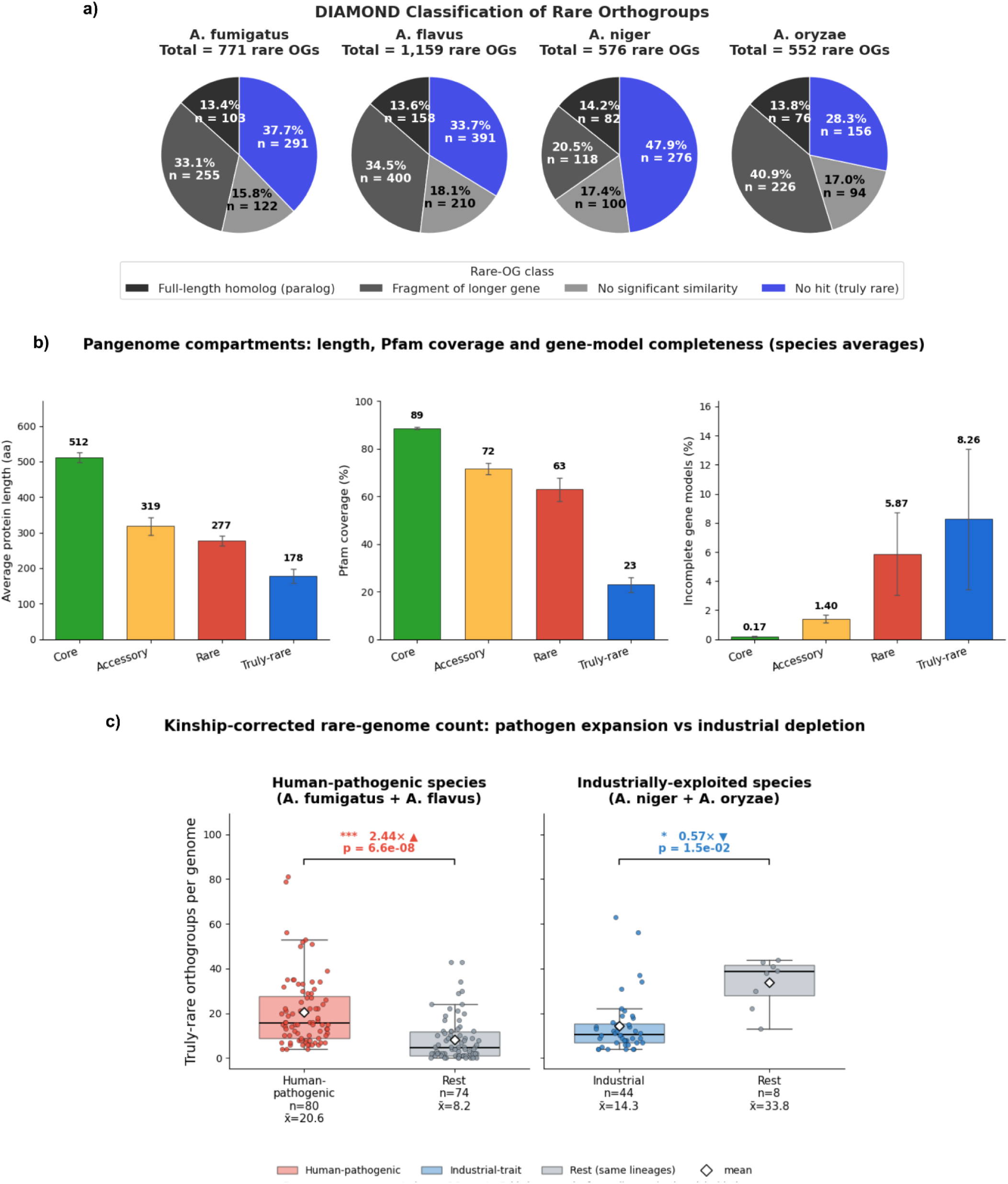
Reclassification of the rare genome. a) DIAMOND classification of rare orthogroups (OGs). Each rare OG was BLASTed against its own species’ Core and Accessory protein pool and assigned to a full-length homolog (paralog), fragment of a longer gene, no-significant-similarity, or No-hit (truly rare and species specific). b) Protein length, Pfam coverage and gene-model completeness across the four Core, Accessory, Rare, and Truly Rare genome, averaged across the four species (bars = species means, error bars = SD). c) Per-genome counts of truly-rare orthogroups in the pooled human-pathogenic species (A. fumigatus + A. flavus) and pooled industrially-exploited species (A. niger + A. oryzae), with means shown as white diamonds. Fold-changes and p-values come from a linear mixed model with the genome-wide kinship matrix as random-effect covariance and species as a fixed effect.

The median predicted protein length was 419 to 442 aa for core OGs, 222 to 277 aa for accessory, 171 to 188 aa for the rare, and 122 to 151 aa for the truly-rare genome. To confirm that the reduced length in the rare genome reflects genuinely short genes rather than truncated genes, we applied a reference-free ORF completeness check (marked incomplete gene if it lacks an ATG start codon or a terminal stop codon, has a CDS length not divisible by three, or lays within a conservative 100 bp of a contig edge), the first three criteria following the NCBI definition of a partial coding sequence [49]. While the Core genome showed a 0.17% truncation, the Accessory genome showed a 1.40% truncation, and the Rare genome 5.87%. The Truly-rare genome showed a 8.26% truncation, meaning that over 90% of truly-rare gene models are complete and their reduced length reflects short genes rather than assembly truncation. What did distinguish the truly-rare genome was its novelty, measured by Pfam annotation rates, where only 19-27% of the truly-rare OGs carry any recognizable Pfam domains, compared with 56-67% for the broader rare genome, 69-74% for accessory, and 88-89% for core genes. All genome averages are illustrated in Figure 6.b.

Targeted NCBI BLAST of high-GC-deviation truly-rare OGs initially flagged two high-confidence cross-kingdom xenologs: an *A. flavus* truly-rare OG (OG0014462) matching a *Stenotrophomonas maltophilia* zonular occludens toxin domain at 98% identity (Expect-value (E) = 0) and an *A. fumigatus* truly-rare OG (OG0010578) matching the tick *Rhipicephalus sanguineus* at 96% identity (E = 1 × 10⁻²³). Both xenologs sat in the No-hit DIAMOND class and survived the truly-rare filter. On closer inspection, however, neither could be validated as genuine horizontal transfers. OG0010578 was matched to a low-complexity Ser-Gly-Tyr repeat protein, while OG0014462 was embedded in a mobile element-like cluster, a prophage/ plasmid-associated cassette. We therefore flagged the contig as contaminated and removed all contaminant OGs from subsequent rare-genome analyses. In the end, neither of the two cross-kingdom transfer candidates survived the validation, and instead marked contamination.

The per-genome count of truly-rare OGs varies between phenotype groups, and also across species. Per-genome rare and truly-rare OG counts, with phenotype and species labels, are given in Table S4. As rare-gene count can be heritable and therefore impacted by phylogenetic distance, all contrasts were additionally tested with genome-wide kinship matrix (GRM) as random-effect covariance and species as a fixed effect to confirm the phenotype effect is not simply a phylogenetic effect. Modelling the two human-pathogenic species (A. fumigatus and A. flavus) together in a linear mixed-effects model with species as a random effect, a clear 2.44-fold expansion (p = 6.6 × 10⁻⁸, kinship corrected) of truly-rare OGs per genome than the rest was measured. The opposite trend was observed in the industrial species (A. niger and A. oryzae), where industrial-trait strains carried 0.57-fold (p = 0.015, kinship corrected) as many truly-rare OGs per genome than the rest (Figure 6.c). Together, a clear bidirectional signal is revealed, with human-pathogenic strains carrying an average of 12.4 more rare OGs and industrial strains carrying an average of 19.5 fewer rare OGs per genome.

The rare genome is the only layer in which the two lifestyles separate reproducibly and in a bidirectional way.

### 3.7 Asymmetric population structure within and between species

To confirm that whole-genome structure recovers the four species, and to justify the kinship correction applied throughout the association tests, we assessed population structure in two ways: Mash-based whole-genome clustering, and how closely each phenotype follows the phylogeny.

Mash-based whole-genome clustering was conducted first. Optimal k were determined by silhouette/ elbow criteria, and was k = 2 for *A. fumigatus* (cluster sizes 82, 6), k = 5 for *A. flavus* (sizes 25, 17, 13, 9, 6), k = 3 for *A. niger* (sizes 14, 3, 2) and k = 3 for *A. oryzae* (sizes 21, 7, 5). As shown in Figure 7, the four-species combined Mash distance matrix resolved into three rather than four whole-genome blocks. *A. fumigatus* and *A. niger* form their own clean clusters at the expected inter-species Mash separation, but *A. flavus* and *A. oryzae* merge into a single cluster indistinguishable from within.

**Figure 7.**
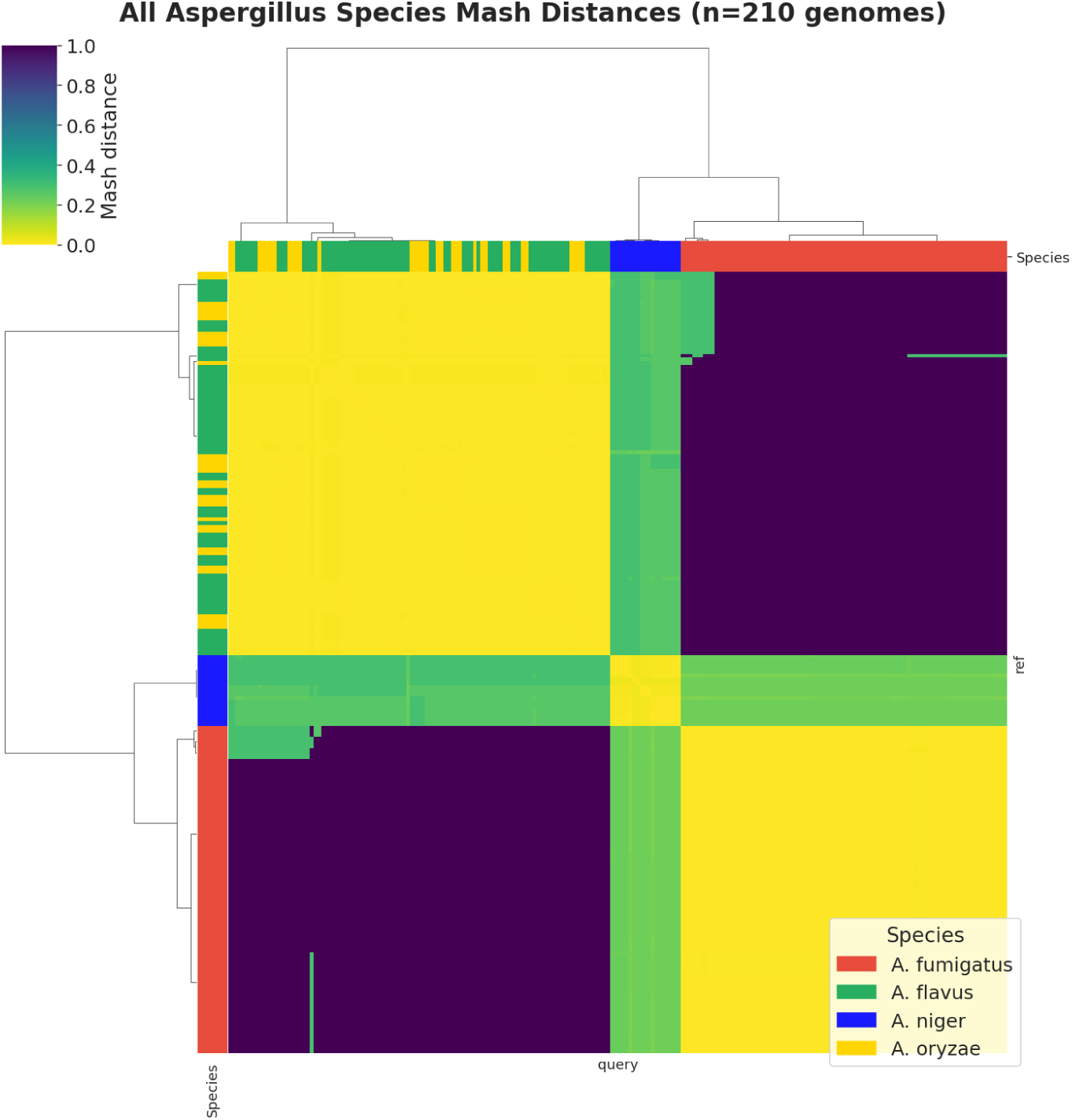
All-species Mash whole-genome distance clustering. Hierarchically clustered heatmap (Ward linkage) of pairwise Mash k-mer distances among all 210 QC-passed genomes. A. flavus and A. oryzae merge into one shared low-distance block.

We then measured how closely each phenotype follows the phylogeny, using the Fritz & Purvis D statistic on the recombination-corrected per-species trees. D near 0 indicates a trait distributed as expected under inheritance, and D near 1 indicates random scattering across the tree. Every testable phenotype was significantly clustered relative to a random null (p ≤ 0.007), and none differed from a Brownian expectation. Clustering was strongest in *A. fumigatus* (D = 0.005 environmental, D = 0.016 human-pathogenic), where clinical and environmental isolates occupy separate parts of the tree rather than being interspersed. In *A. flavus*, human pathogenicity was far less constrained by phylogeny (D = 0.279) than plant pathogenicity (D = −0.094), the most clustered phenotype in the dataset. *A. niger* gave D = 0.212 for industrial and D = −0.016 for environmental strains. *A. flavus* environmental strains were too few to test reliably (n = 4, p = 0.054), and *A. oryzae* was not testable under its 32:1 split.

In *A. fumigatus*, phenotype is nearly inseparable from position on the tree (D = 0.005-0.016), and every other testable phenotype shows the same pattern in weaker form. Both the strongly asymmetric within-species structure, most notably the 82:6 split in *A. fumigatus*, and the collapse of *A. flavus* and *A. oryzae* into one block therefore confirms that background relatedness is substantial at every level of this dataset, and justify modelling and correcting for it using the kinship matrix as random-effect covariance throughout the association tests.

## 4. DISCUSSION / CONCLUSION

The analysis of four *Aspergillus* species reveals how these fungi adapt to different environments through changes in their species-specific genes. We found a clear bidirectional pattern: human-pathogenic strains expand their species-specific gene repertoires, while industrially domesticated strains contract them. This pattern appears independently in two separate species pairs. Human-pathogenic strains show a 2.44-fold expansion of truly-rare orthogroups per genome in the pooled *A. fumigatus* and *A. flavus* model (p = 6.6 × 10⁻⁸, kinship corrected), while industrial strains of *A. niger* and *A. oryzae* carry 0.57-fold as many (p = 0.015, kinship corrected).

The absence of convergent evolution across species is of interest. Past comparative genomics of the *Aspergillus* genus has shown both a shared metabolic and regulatory backbone [50], [51] and species-specific gene content [48], [52], [53], [54], [55]. However, no study has formally tested whether *A. fumigatus* and *A. flavus* pathogenic strains share the same gene-content changes for the same phenotype. When we tested this directly, we found no evidence of shared evolutionary solutions. 448 orthogroups were tested in both *A. fumigatus* and *A. flavus*, where none reached significance in both. The direction of effects for both PAV and CNV results agreed 49% of the time, which is no better than chance (exact binomial p = 0.81 and p = 0.74). The size of the effects were also uncorrelated between species (Spearman ρ = −0.023 and −0.020). We also tested for functional convergence. We examined whether species shared similar biological functions across gene categories and species pairs, but none survived multiple-testing correction (0 of 30 significant at q < 0.05). This means that while both *A. fumigatus* and *A. flavus* become human pathogens, this work predicts that they do so through different genetic mechanisms. The equivalent test for *A. niger* and *A. oryzae*, however, was not estimable, as *A. oryzae* yielded no significant orthogroups under its case/control imbalance.

A complementary check in the Core genome reinforces the conclusion. In fact, the lifestyle signal detected in the species-specific layer cannot be recovered from the literature-curated verified trait genes. 84% of the 128 experimentally validated trait genes are core in all four species, compared with only 41% of background OGs (Mann-Whitney p = 2 × 10⁻²²). None of the 73 virulence genes are strictly pathogen-specific, that is, core in both *A. fumigatus* and *A. flavus* and absent from both *A. niger* and *A. oryzae*. The pathogenic *A. flavus* and its GRAS domesticate *A. oryzae* demonstrate directly that virulence genes cannot by themselves separate pathogens from domesticates. All 8 experimentally validated core *A. flavus* virulence genes are also core in *A. oryzae*, including the aflatoxin pathway regulators aflR and aflS.

Gene content alone therefore cannot explain why one species produces aflatoxin and the other is food-grade, and for the aflatoxin cluster the explanation has already been established directly. Most industrial strains of *A. oryzae* retain the cluster in an inactive state, carrying two substitutions in the aflR promoter that hinder productive aflR expression [56], and amino acid substitutions that inactivate aflS (aflJ) [57]. Both aflatoxin regulators are present but neither functional. *A. oryzae*’s food-grade status would therefore reflect gene regulatory inactivation rather than gene loss.

A methodological insight emerged from our analysis. The rare gene category traditionally used in pangenome studies contains substantial noise, where 35-55% consists of gene family duplicates and fragmented genes rather than truly strain-exclusive within the species’ genes. We hence filtered out this noise and focused only on genes with no similarity to their own species’ core and accessory proteins. The bidirectional pattern remained significant (industrial p = 0.015; pathogenic p = 6.6 × 10⁻⁸) but was more biologically interpretable. A comparable pattern was recently reported for the *Escherichia coli* pangenome, where rare genes were predominantly fragments and divergent copies of core or accessory genes [58], suggesting this feature is shared across bacterial and fungal pangenomes.

In *A. fumigatus*, the rare-gene expansion is concentrated in a single dominant clinical cluster (Mash k = 2, cluster sizes 82 and 6), consistent with the structured global *A. fumigatus* population previously described. Two candidate cross-kingdom xenologs initially recovered from the truly rare genome did not survive validation: one proved to be a low-complexity repeat protein and the other sat within a mobile-element cassette on a contig we subsequently flagged as contaminated. Both were removed. This illustrates a general hazard of rare-genome analysis, in which the same properties that mark a gene as genuinely novel, being no internal homolog and deviant GC content, can also reveal itself to be a contaminant, and reinforces the necessity of explicit contig-level validation.

Comparatively, *A. flavus* showed no such concentration. Its clinical isolates are instead spread across five independent within-species clades (Mash k = 5). Such results would be consistent with the view of *A. flavus* human pathogenic strains as opportunistic, exploiting an existing plant-pathogen toolkit rather than host-specialised adaptation [59], [60]. The one secondary-metabolite cluster tracking human pathogenicity in *A. flavus* (FAM_00077) was gained in human-pathogenic strains but lost in plant-pathogenic ones, following a trend of highly variable secondary metabolite cluster repertoire across the species [61]. These findings suggest that *A. fumigatus* causes aspergillosis through acquisition of tissue-invasion modules, whereas *A. flavus* redirects its pre-existing agricultural-pathogenic toolkit in an opportunistic manner.

In parallel, *A. niger* and *A. oryzae* show different patterns under industrial domestication. Both species are industrial cell factories, *A. oryzae* for fermented foods and *A. niger* for enzymes and organic acid production. *A. oryzae* is also understood to be a domesticated derivative of *A. flavus*, selected over the past several thousand years [48]. The proximity of *A. oryzae* to *A. flavus* is reflected in our Mash analysis, where the two species merge into a single whole-genome k-mer cluster while *A. niger* forms its own cluster. This is consistent with their ∼99.5% genomic identity and lack of separation by ANI clustering [62], although finer-scale SNP analyses distinguish the two [63]. Together, these results suggest that under industrial selection the rare genome contracts rather than expands. Pooled across the two industrially exploited species, industrial-trait strains carry a mean of 14.3 truly-rare orthogroups per genome against 33.8 in their non-industrial counterparts, a difference of 19.5 orthogroups. *A. niger* industrial strains are additionally strongly enriched for the PDR/CDR ABC-transporter efflux family (KEGG_TC 3.A.1.205, q = 0.016), which could be a result of bioreactor selection for xenobiotic and byproduct efflux [52].

This means that industrial domestication and human pathogenicity act on the same rare genome of the pangenome but in opposite directions. Pathogenic species such as *A. fumigatus* and *A. flavus* expand rare gene content under host selection, while industrial species such as *A. niger* and *A. oryzae* lose rare gene content through the bottlenecked selective cleanses that produced the modern industrial strains. The direction of each shift fits the niche. A pathogen occupies an open and variable niche, thus expands and accumulates functions to meet unpredictable host stresses. An industrial strain is a specialist held in controlled production conditions, and therefore adapts and simplifies toward a smaller set of useful functions. The same streamlining is seen under adaptive laboratory evolution (ALE), where laboratory evolution under controlled conditions similarly reduces an organism’s systems toward a subset of beneficial functions [64].

The clinical and biotechnological implications are clear. Virulence genes identified in *A. fumigatus* will not necessarily apply to *A. flavus* as each species reaches pathogenicity through its own evolutionary route. Pan-*Aspergillus* biomarkers will be difficult to develop. Instead, species-specific markers derived from each species-specific pangenome gene category offer a more realistic target. For *A. fumigatus*, the truly-rare genome provides concrete candidates for clinical biomarker development, although the low Pfam annotation rate (19-27%) means most such candidates remain functionally uncharacterised. In biotechnology, *A. niger* and *A. oryzae* industrial strains show contracted species-specific pangenome gene categories, with *A. niger* ABC-transporter efflux systems enriched, which is where domestication has acted and where engineering efforts to modify strain capabilities could most usefully be focused.

## 5. LIMITATIONS

### 5.1 Sample size heterogeneity and statistical power constraints

The dataset shows substantial sample size imbalance across species. *A. niger* (n = 19) and *A. oryzae* (n = 33) are approximately 2.5- 4.5 times smaller than *A. fumigatus* (n = 88) and *A. flavus* (n = 70). This imbalance creates species-specific power limitations that can affect the statistical framework. For *A. oryzae*, the extreme 32:1 industrial to rest partition violates the minimum case-control ratio requirements for stable linear mixed model estimation, making its species-level pan-GWAS contrast and convergence analysis untestable. Similarly, *A. niger* industrial-vs-rest contrasts suffer from reduced statistical power, with minimum detectable odds ratios of 22.33 at 80% power. Therefore, the lack of significant *A. niger* industrial pan-GWAS associations (2 PAV, 1 CNV) cannot be interpreted as evidence for absence of effect. Fortunately, these single-species power limitations do not compromise the bidirectional hypothesis, as the pooled cross-species mixed-effects analysis accounts for genome-size heterogeneity into random effects and allows both directional effects to be assessed with sufficient statistical power.

### 5.2 Population representation and clinical isolate redundancy

The *A. fumigatus* genomic dataset includes substantial representation from COVID-19-associated pulmonary aspergillosis (CAPA) outbreak investigations (BioProjects PRJNA673120, PRJNA697844), introducing potential population structure artifacts through sampling of closely related clinical isolates. Mash distance-based clustering reveals asymmetric population structure (cluster sizes: 82, 6), suggesting within-cluster genetic redundancy that may inflate effective sample sizes for association testing. Phylogenetic signal analysis shows the same problem in a stronger form: phenotype in this species is almost perfectly confounded with phylogeny (D = 0.005-0.016), with clinical and environmental isolates occupying distinct regions of the tree rather than being interspersed. Kinship-corrected LMM implementation partially mitigated this concern by using the full genome-wide relatedness matrix as a random effect, absorbing background population structure. However, where phenotype and ancestry are this closely aligned, correction cannot fully separate the two, and the associations reported for *A. fumigatus* should be read as a lower bound on the recoverable signal.

### 5.3 Phenotype classification methodology and label uncertainty

Within the pan-GWAS analyses, human error in the manual phenotype curation can potentially introduce label misclassification noise that could bias association testing. Systematic evaluation under strict confidence thresholds needs to be performed.

## 6. DATA AND CODE AVAILABILITY

All raw genome assemblies are publicly available through NCBI under the accessions listed in qc_passed_metadata_enriched.csv. The complete FunPan pipeline (README.md, Code/), including all shell scripts, the conda environment specification (funpan), and per-species reproducible analysis notebooks (NB0_DataPrep.ipynb, NB1_Pangenome.ipynb, NB2_PanGWAS.ipynb, NB3_Convergence.ipynb, NB4_RareGenome.ipynb, NB5_CoreGenome.ipynb) and analysis modules (funpan_utils.py, funpan_classify.py, funpan_pangenome.py, funpan_gwas.py, funpan_convergence.py, funpan_phylo.py) will be deposited in Zenodo upon publication. Per-species result files (PAV / CNV / BGC / GCF matrices, kinship and SNP-PC tables, OG consensus annotations, pan-GWAS association tables, convergence outputs, and rare-genome characterisation outputs) are organised under NB0_Results/, NB1_Results/, NB2_Results/, NB3_Results/, NB4_Results/, and NB5_Results/.

## ACKNOWLEDGEMENTS

We would like to thank Jonathan Monk and Pablo Cruz-Morales for the helpful technical discussions. This work was funded by The Novo Nordisk Foundation (Grant Number NNF24SA0100980).

## Author Contributions

M.K., P.P., and E.K. designed research; M.K., P.P., O.A., and E.K. contributed analytical methods and tools, M.K. acquired data and performed analysis; M.K. drafted the manuscript and figures; M.K., O.A., and P.P. carried out critical revision and final manuscript approval.

## 7. SUPPLEMENTARY MATERIALS

*Supplementary Table 1. Per-genome assembly metrics, isolation metadata, and phenotype classification for the 210 quality-filtered Aspergillus genomes. Each row corresponds to one genome. BUSCO completeness, contig count and contig N50 are the quality-control metrics used for genome filtering. The Provenance_keywords and Phenotype_keywords columns list the regex patterns that triggered each assigned label, and the Provenance_fields and Phenotype_fields columns list the metadata fields in which those patterns were found. The Negations_applied column records any negation rule that suppressed a candidate label. The Used_in_panGWAS column marks the 206 genomes with a defined phenotype that entered association testing*.

*Supplementary Table 2. Literature-curated virulence and industrial trait genes and their pangenome class across four Aspergillus species. Each row is one experimentally validated gene-trait assignment covering 123 genes, five of which were validated in two species and appear twice. Characterised_in is the species the trait was experimentally demonstrated in. Genus_orthogroup is the combined four-species orthogroup the reference protein anchors to. Class columns give whether the orthogroup falls in the core, accessory, or rare genome of each species, or is absent from it. PMID and Reference give the supporting publication*.

*Supplementary Table 3. DIAMOND reclassification of every rare orthogroup against its own species’ core and accessory protein pool. Each row corresponds to one rare orthogroup. Full-length homolog denotes a paralog of a non-rare gene (≥30% identity with ≥80% query and subject coverage), Fragment of longer gene denotes high subject coverage with low query coverage, No significant similarity denotes a DIAMOND hit below threshold, and No hit denotes no alignment to the core or accessory pool at all. Identity_pct, Query_coverage_pct and Subject_coverage_pct give the alignment statistics of the best hit, and Best_hit_protein its identifier. Truly_rare marks the No-hit orthogroups retained for all subsequent rare-genome analyses*.

*Supplementary Table 4. Per-genome rare and truly-rare orthogroup counts underlying the kinship-corrected burden model. Each row corresponds to one genome. Rare_OGs is the number of rare-class orthogroups the genome carries, and TrulyRare_OGs the subset assigned to the No-hit DIAMOND class. Lineage_pool indicates which of the two pooled cross-species models the genome contributed to, and Model_group its case or control assignment within that model. The four genomes with an Unknown phenotype are excluded*.

